# Mutations in *MYLPF* cause a novel segmental amyoplasia that manifests as distal arthrogryposis

**DOI:** 10.1101/2020.05.06.071555

**Authors:** Jessica X. Chong, Jared C. Talbot, Emily M. Teets, Samantha Previs, Brit L. Martin, Kathryn M. Shively, Colby T. Marvin, Arthur S. Aylsworth, Reem Saadeh-Haddad, Ulrich A. Schatz, Francesca Inzana, Tawfeg Ben-Omran, Fatima Almusafri, Mariam Al-Mulla, Kati J. Buckingham, Tamar Harel, Hagar Mor-Shaked, Periyasamy Radhakrishnan, Katta M Girisha, Shalini S. Nayak, Anju Shukla, Klaus Dieterich, Julien Faure, John Rendu, Yline Capri, Xenia Latypova, Deborah A. Nickerson, David Warshaw, Paul M. Janssen, University of Washington Center for Mendelian Genomics, Sharon L. Amacher, Michael J. Bamshad

## Abstract

We identified ten persons in six consanguineous families with Distal Arthrogryposis (DA) who had congenital contractures, scoliosis, and short stature. Exome sequencing revealed that each affected person was homozygous for one of two different rare variants (c.470G>T, p.(Cys157Phe) or c.469T>C, p.(Cys157Arg)) affecting the same residue of *myosin light chain, phosphorylatable, fast skeletal muscle* (*MYLPF)*. In a seventh family, a c.487G>A, p.(Gly163Ser) variant in *MYLPF* arose *de novo* in a father, who transmitted it to his son. In an eighth family comprised of seven individuals with dominantly-inherited DA, a c.98C>T, p.(Ala33Val) variant segregated in all four persons tested. Variants in *MYLPF* underlie both dominant and recessively inherited DA. Mylpf protein models suggest that the residues associated with dominant DA interact with myosin whereas the residues altered in families with recessive DA only indirectly impair this interaction. Pathological and histological exam of a foot amputated from an affected child revealed complete absence of skeletal muscle (i.e., segmental amyoplasia). To investigate the mechanism for this finding, we generated an animal model for partial MYLPF impairment by knocking out zebrafish *mylpfa*. The *mylpfa* mutant had reduced trunk contractile force and complete pectoral fin paralysis, demonstrating that *mylpf* impairment most severely affects limb movement. *mylpfa* mutant muscle weakness was most pronounced in an appendicular muscle and was explained by reduced myosin activity and fiber degeneration. Collectively, our findings demonstrate that partial loss of MYLPF function can lead to congenital contractures, likely as a result of degeneration of skeletal muscle in the distal limb.

## Introduction

The distal arthrogryposes (DA) are a group of Mendelian conditions with overlapping phenotypic characteristics, shared genetic etiologies, and similar pathogenesis.^1^ Clinically, the DAs are characterized by non-progressive congenital contractures of the limbs, most commonly affecting the hands, wrists, feet, and ankles. Congenital contractures of the face, ocular muscles, neck webbing, pterygia, short stature, and scoliosis are less frequent, variable findings that facilitate delineation among the most common DA conditions: DA1^2^ (MIM 108120), DA2A^3^ (Freeman-Sheldon syndrome [MIM 193700]) and DA2B^4^ (Sheldon-Hall syndrome [601680]). Variants in any one of sixteen different genes can underlie DA but the overwhelming majority of known pathogenic variants occur in just five genes (*TPM2* (MIM 190990), *TNNI2* (MIM 191043), *TNNT3* (MIM 600692), *MYH3* (MIM 160720), *MYH8* (MIM 160741)).^5,6^ Yet, collectively pathogenic variants are identified in only ∼60% of families diagnosed with a DA, so the precise genetic etiology remains unknown in nearly half of DA families.

Most of the genes that underlie DA encode sarcomeric components of skeletal muscle fibers. Thus, genes encoding sarcomeric proteins have long been considered priority candidates for DA. Sarcomeres are the fundamental contractile structure of muscle, wherein myosin-rich thick filaments interact with actin-based thin filaments to generate contractile force. Each skeletal muscle myosin heavy chain protein (MyHC) has two distinct light chain proteins bound to the myosin lever arm: an essential light chain that is nearest to the myosin head, and a regulatory light chain protein that can be phosphorylated and is located closer to the myosin tail region.^7^ Light chain proteins are needed to stabilize the myosin lever arm so that myosin can generate maximum force and velocity as revealed by in vitro studies of isolated myosin extracts deficient in light chain proteins.^8,9^

To discover novel genes underlying DA, we performed exome sequencing (ES) on 172 families in which pathological variants in genes known to underlie DA1, DA2A, and DA2B had not been identified via Sanger sequencing. We identified putative pathogenic variants in 80 (47%) of these families including 44 families with mutations in 20 genes not known to underlie DA. Affected individuals from two families, including an affected child (Family B) who had complete absence of skeletal muscle (i.e., segmental amyoplasia) in a foot (Figure 1), were each homozygous for the same variant (c.470G>T) in the gene, *MYLPF*, which encodes the fast-type skeletal muscle regulatory light chain. Data sharing via MatchMaker Exchange (MME) and directly with commercial diagnostic labs identified six additional families with similar phenotypic features and rare variants in *MYLPF*, including two families in which the condition was transmitted from parent to offspring (Table 1, Families G and H). The mouse *Mylpf* knockout mutant is born without skeletal muscle and dies soon after birth because of respiratory failure,^10^ suggesting that human pathogenic *MYLPF* variants are likely to be hypomorphic alleles. To test this hypothesis, we knocked out the more prominent of the two zebrafish *mylpf* genes, *mylpfa*, and characterized development and function of *mylpfa* mutant skeletal muscle.

**Table 1.**
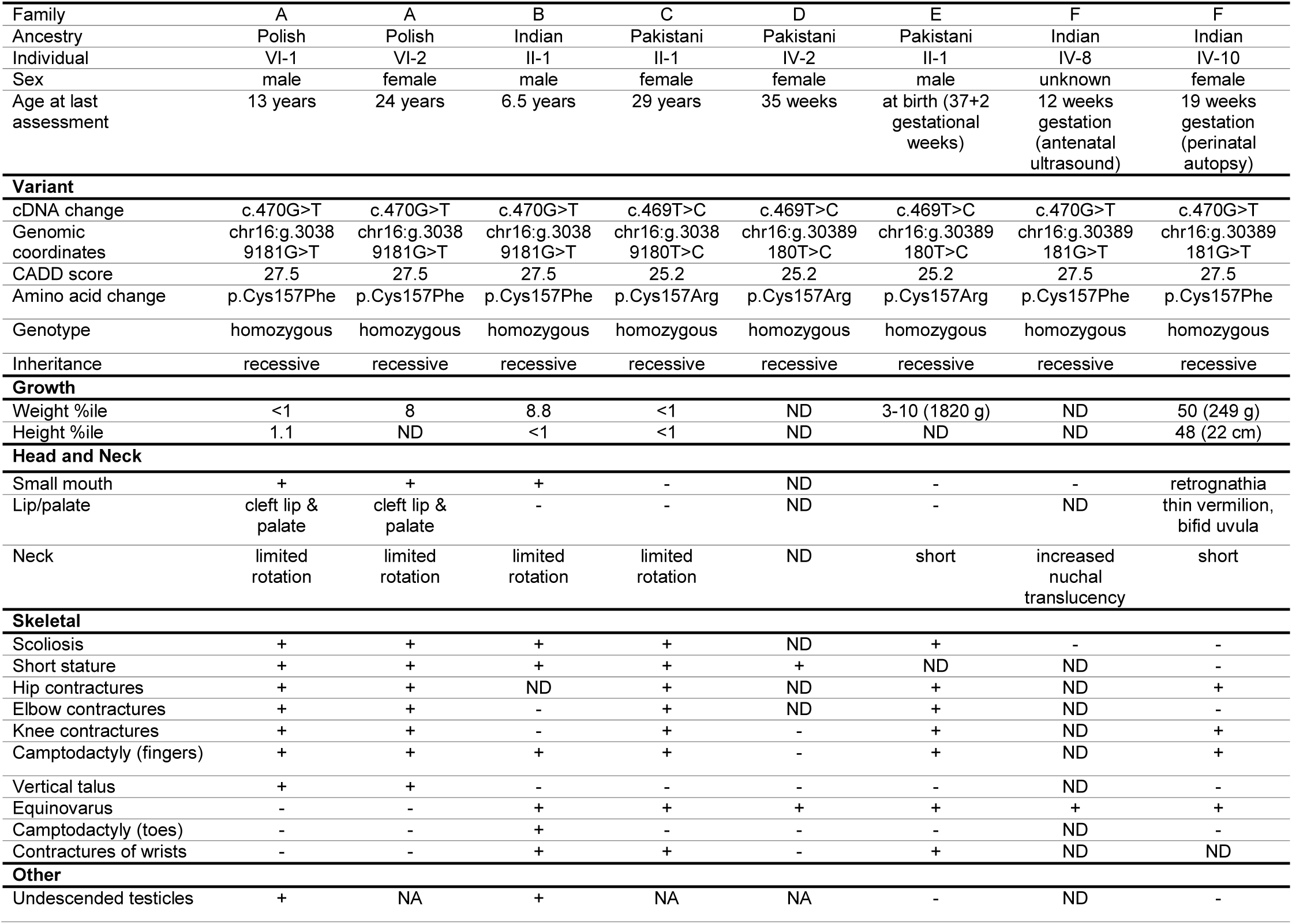
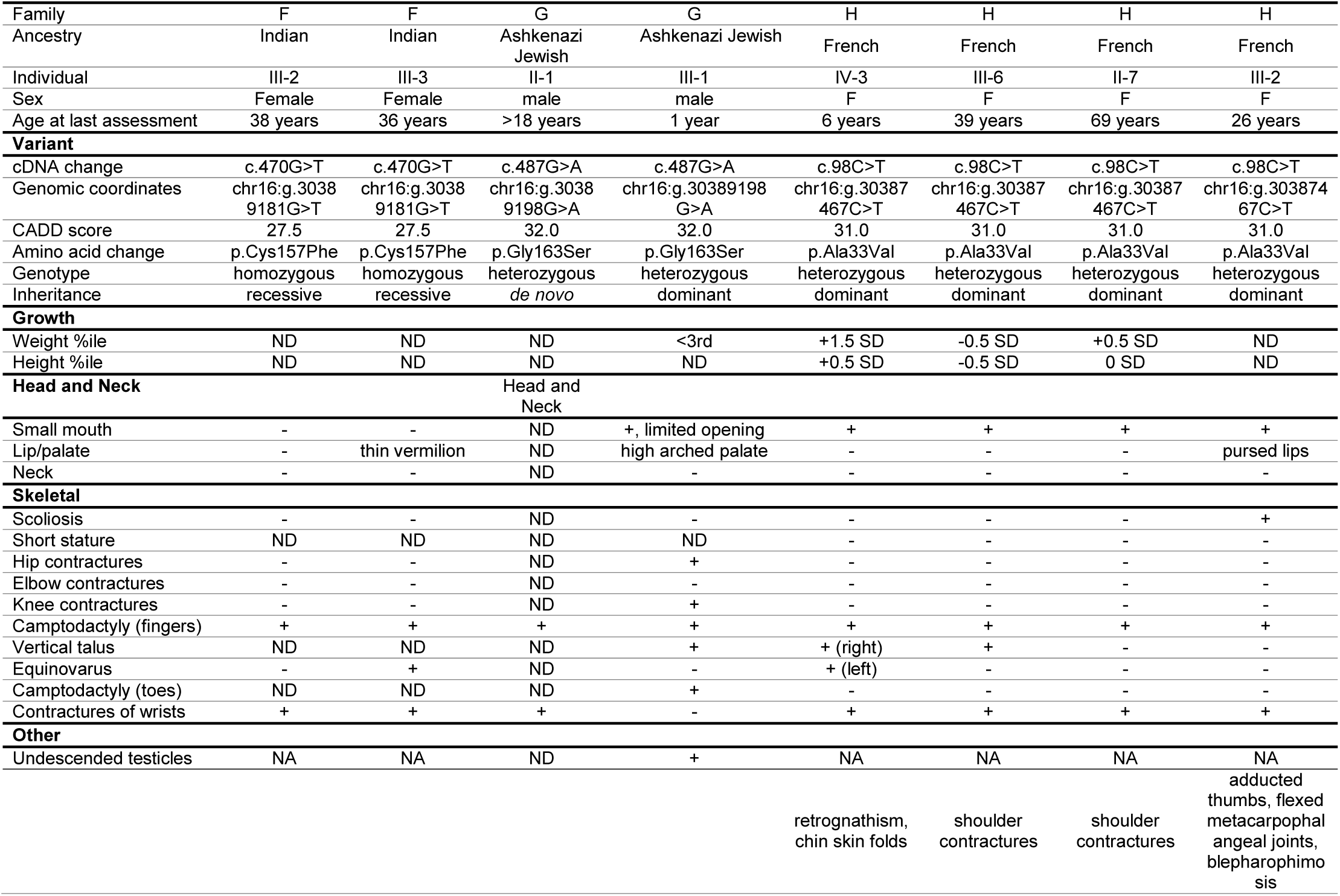
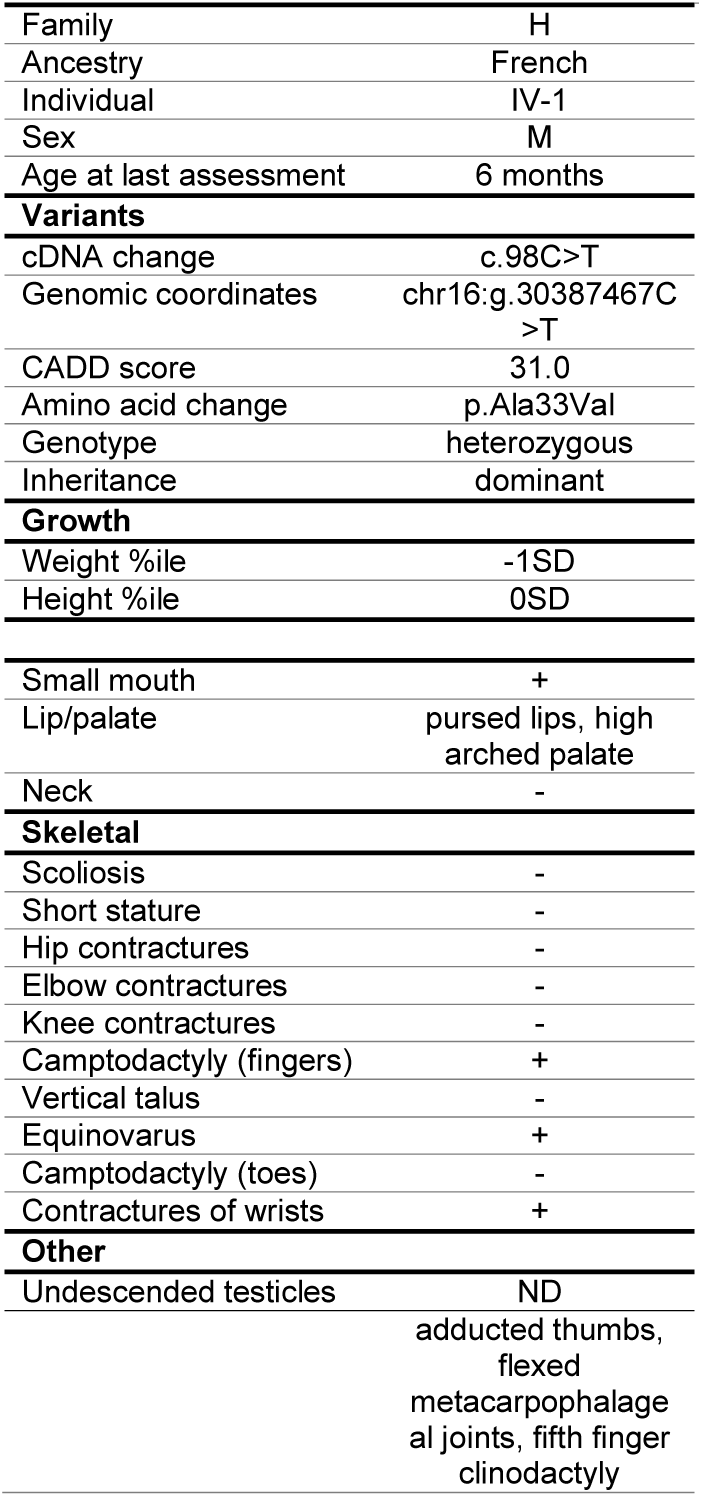
Mutations and Clinical Findings of Individuals with Distal Arthrogryposis type 1 due to variants in *MYLPF*. This table provides a summary of clinical features of affected individuals from families in which mutations in *MYLPF* were identified. Clinical characteristics listed in the table are primarily features that delineate DA1. Plus (+) indicates presence of a finding, minus (-) indicates absence of a finding. * = described per report. ND = no data were available. NA = not applicable. CADD = Combined Annotation Dependent Depletion v1.6. cDNA positions named using HGVS notation and RefSeq transcript NM_002470.3. Predicted amino acid changes are shown.

**Figure 1:**
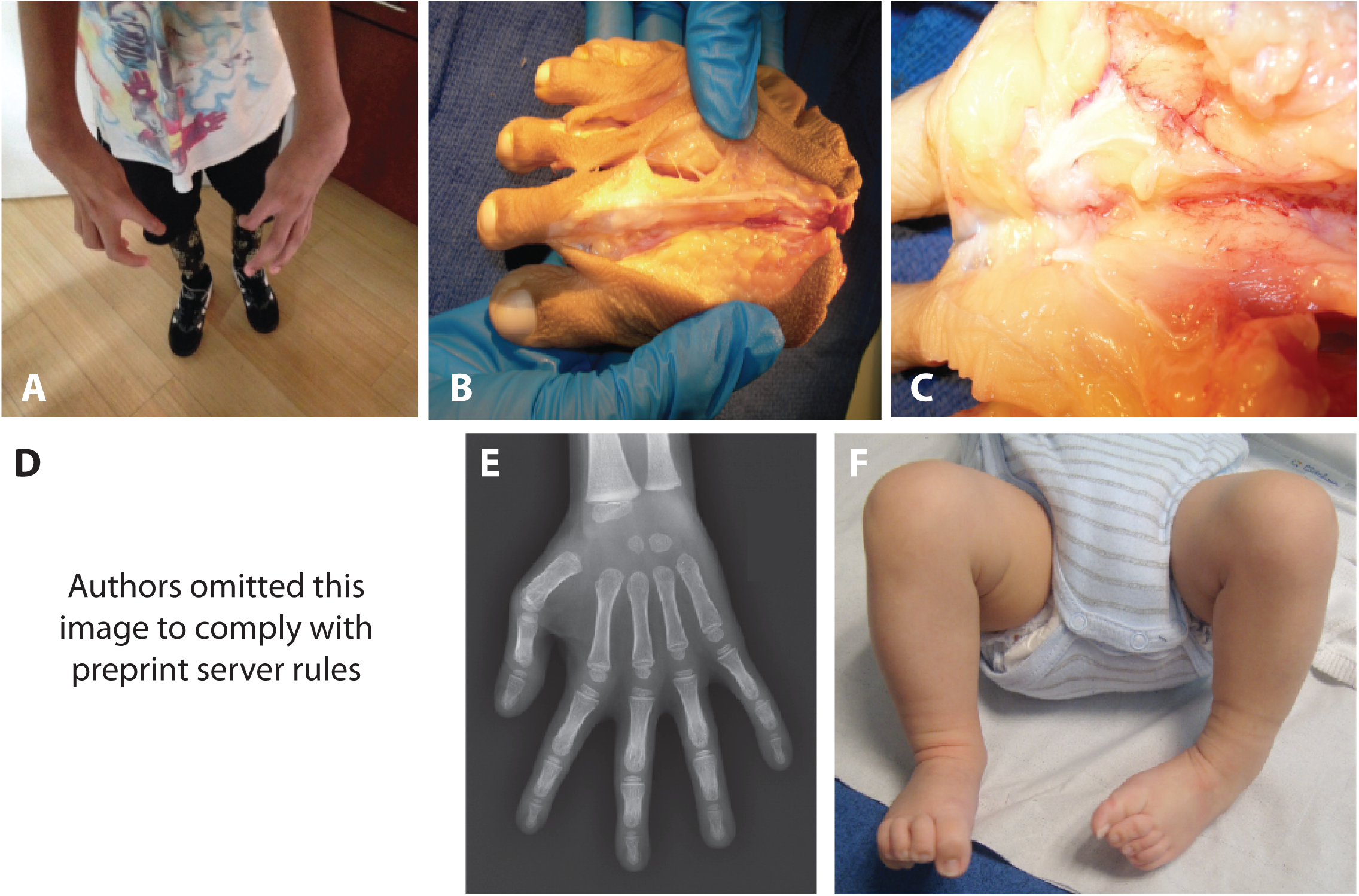
Phenotypic Characteristics of Individuals with Recessive or Dominant Distal Arthrogryposis type 1 due to variants in *MYLPF*. (**A-C**) Characteristics shown in Family B II-1 with recessive DA1 A) camptodactyly of the fingers and radial deviation of the wrists. B,C) Gross pathology of the right foot illustrating absence of skeletal muscles. **(D-F**) Characteristics in Family H IV-1 with dominant DA1 D) [authors removed this image to comply with preprint server rules] pursed lips, camptodactyly of the fingers, adducted thumbs E) clinodactyly of the fifth digit F) bilateral clubfoot. Table 1 contains a detailed description of the phenotype of each affected individual and Figure S1 provides a pedigree of each family with DA1 due to variants in *MYLPF*.

Zebrafish are a well-established model for investigating muscle structure, development, and disease mechanisms.^11–14^ Zebrafish rapidly generate functional myofibers that produce both spontaneous and evoked contractions at one day post-fertilization (dpf). By this stage, muscle fiber type is also readily apparent, with fast-twitch muscles identified by expression of *mylpfa* and other markers.^15–18^ Between 1 and 3 dpf, muscle precursors migrate away from their origin to produce new *mylpfa-*positive muscles, including fin muscles and the posterior hypaxial muscle.^19–23^ Herein, we show that partial loss of *Mylpf* function in zebrafish can recapitulate DA1, demonstrate that variants in *MYLPF* underlie DA1, and use a zebrafish model to provide explanations for the limb muscle loss observed in DA1 due to pathogenic variants in *MYLPF*.

## Materials and Methods

### Discovery Cohort

From our cohort of 463 families (1,582 individuals) with multiple congenital contractures, we selected 172 families in which pathological variants had not been identified. All studies were approved by the institutional review boards of the University of Washington and Seattle Children’s Hospital and informed consent was obtained from each participant or their parents.

### Exome Sequencing and Variant Analysis

ES was performed by the University of Washington Center for Mendelian Genomics as described previously.^24^ In brief, data were annotated with the Variant Effect Predictor v89^25^ and analyzed using GEMINI 0.20.2.^26^ Variants unlikely to impact protein-coding sequence (for which GEMINI impact_severity = LOW), variants flagged by the Genome Analysis Toolkit (GATK) as low quality, and variants with an alternative allele frequency > 0.005 in any super-population in EVS/ESP6500, 1000 Genomes (phase 3 release), or the gnomAD Browser (v.2.0.1) were excluded. Variants that were frequent (alternative allele frequency >0.5) in an internal database of >7,500 individuals (Geno2MP v1.7 release) were excluded. Candidate genes were identified by filtering under these parameters for variants matching the predicted pattern of inheritance (i.e. autosomal *de novo*, homozygous recessive, compound heterozygous, X-linked *de novo*, X-linked recessive, and X-linked dominant models).

### Fish maintenance and husbandry, transgenes, and mutant construction

All animal protocols used in this study are approved by the Institutional Animal Care and Use Committees at The Ohio State University, the University of Vermont, and the University of Maine. Standard practices were used for zebrafish husbandry and maintenance.^27^ Transgenic and mutant zebrafish strains were maintained on the AB wild-type background. Transgenic lines used in this study are *Tg(mylpfa:lyn-Cyan)fb122*,^28^ *Tg(myog:Hsa*.*HIST1H2BJ-mRFP)fb121* (abbreviated *myog:H2B-mRFP*),^29^ and *Tg(smyhc1:EGFP)i104*,^30^ which are combined in a ‘3MuscleGlow’ triple-transgenic line.^31^ The two *mylpfa* mutant lines described in this study were generated using established CRISPR/Cas9 protocols.^32^ One-cell embryos were injected with Cas9 mRNA and guide RNA targeting Exon 2 (5’-TTGAGGCCAACACGTCCCTA-3’) or Exon 3 (5’-GGTGAAGTTGATTGGGCCGC-3’), raised to adulthood, and outcrossed. F1 progeny were screened using HRMA to identify founders carrying CRISPR-induced *mylpfa*^*oz43*^ and *mylpfa*^*oz30*^ lesions. All lesions were sequence confirmed in homozygotes. Mutations were outcrossed at least two generations after CRISPR injection before phenotypic analyses.

### Zebrafish Immunohistochemistry and RNA in situ hybridization

Embryos and larvae carrying the *Tg(smyhc1:EGFP)i104* transgene were fixed and immunolabeled with Rbfox1l (1:500)^33^ and F310 (1:100, Developmental Studies Hybridoma Bank) antibodies. RNA in situ hybridization was performed as described^34^ using *mylpfa*^18^ and *mylpfb* riboprobes. For the latter, a 408 bp *mylpfb* fragment was amplified from zebrafish cDNA using forward 5’-AGTGGCCCCATCAACTTTACTG-3’ and reverse 5’-AGCCCAAATGCCAACAAACC-3’ primers and cloned into a PCR4-TOPO vector for subsequent probe synthesis.

### Live imaging of muscle structure

The following transgenes were used for live imaging: *Tg(mylpfa:lyn-Cyan)fb122*^28^ to visualize fast muscle membranes, Tg(*myog:H2B-mRFP)*^29^ to visualize myonuclei, and *Tg(smyhc1:EGFP)i104*^30^ to visualize slow muscle fibers. Time-lapse imaging was performed as described.^23^ For 3.25 dpf (78 hpf) and 4.25 dpf (102 hpf) comparisons, fish were dismounted and raised at 28.5°C between imaging sessions.

### Muscle Contractile Force Measurement

Contractile analysis of 3 dpf larvae was performed as described previously.^35^ Live 3 dpf larvae were anesthetized in 0.02% weight/volume tricaine buffered with Tris·HCl in Krebs-Henseleit solution and mounted on a custom-built set up between a force transducer and a hook. Larvae were stimulated at increasing frequencies and contractile strength measured. The maximal contractile force reached during each contraction was recorded, analyzed, and reported per larva. Fused tetanic contractions occur at 180 Hz. Measurements were compared at each contraction frequency using ANOVA followed by Tukey-Kramer post-hoc comparisons.

### Behavioral analysis

To quantify fin movement, we recorded fish for one full minute, recorded the number of beats on each side of the fish, and then averaged fin movements per side. Escape response was evoked by gentle probing.^36^ Freely moving fish were imaged using a Leica DMC5400 camera mounted on a Leica MZ10F microscope; images were collected in LAS X software and processed in FIJI.

### Assessing Isolated Myosin Molecular Function

Zebrafish carrying *mylpfa*^*oz30*^ and *mylpfa*^*oz43*^ mutations were intercrossed, raised to 2 dpf, and sorted for wild-type or mutant swimming behavior. Proper sorting of mutant versus wild-type sibling embryo genotypes was confirmed on over 100 fish per group. At 4 dpf, fish were prepared, myosin extracted, and the *in vitro* motility assay was performed as described in the Supplementary Methods. Fish were dechorionated, de-yolked, permeabilized, and cut open through the abdomen to expose the de-membranated muscle fibers to subsequent solutions. Two fish larvae were inserted, tail first, into a flow cell constructed from a microscope slide and coverslip, as described previously.^37^ Myosin Extraction Buffer was infused and incubated for one hour, followed by 0.5mg/ml BSA in Actin Buffer and incubated for 2 minutes at 30°C. All solution changes after this point were identical to that previously reported.^38^ In brief, unlabeled actin was infused to effectively eliminate non-functional myosin heads, and then followed by rhodamine-phalloidin labeled actin in Motility Buffer containing ATP. Imaging and actin filament tracking and velocity analysis were conducted as previously described.^39^ Each flow cell contained myosin from two fish, and at least four fields of view were imaged with at least 20 fluorescent actin filaments tracked per field. Velocity of these tracks was averaged across all fields of view for a single statistical N. Experiments for a given group were repeated on 4 flow cells minimum with at least two fully independent replicates on separate imaging days. For illustrative purposes in Figure 3L and Movie S3, actin filaments were tracked using the MTrackJ function in FIJI. Actin filament particles that stayed in the viewing frame throughout 100 frames of imaging were randomly selected and tracked for both wild-type sibling and *mylpfa*^*oz30*^ mutant genotypes.

**Figure 3:**
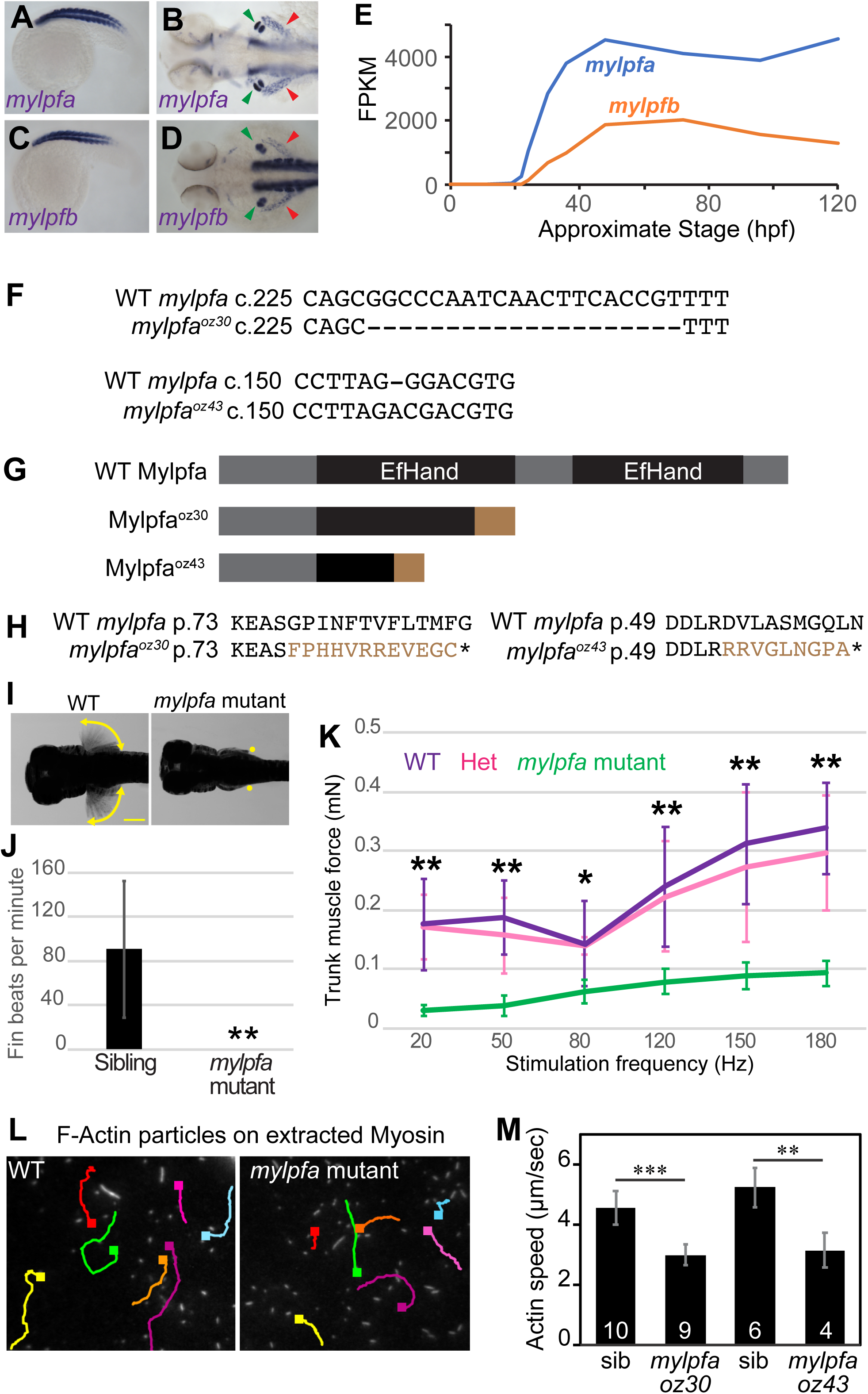
Zebrafish *mylpfa* mutants have weakened myotomes and paralyzed fin muscle. **(A-D)** RNA *in situ* hybridization showing (A, B) *mylpfa* and (C, D) *mylpfb* expression at 20 hpf (A, C) and 52 hpf (B, D). Both genes are expressed exclusively in fast muscle, including somitic muscles, fin muscle (green arrowhead), and posterior hypaxial muscle (red arrowhead). **(E)** Transcript abundance of *mylpfa* (blue) and *mylpfb* (orange) through early larval development, using data provided in the EMBL-EBI expression atlas.^58^ **(F)** Alignment of wild-type (WT) and mutant genomic sequence across the *mylpfa*^*oz30*^ and *mylpfa*^*oz43*^ lesions. *mylpfa*^*oz30*^ is a 20 bp deletion within Exon 3 predicted to frameshift the 169 amino acid protein after amino acid 76, and *mylpfa*^*oz43*^ is a 1 bp deletion within Exon 2 predicted to frameshift the protein after amino acid 52. **(G)** Diagram of wild-type and predicted mutant Mylpfa proteins. Both mutant alleles should truncate the protein within the first EF-hand domain (black boxes) and introduce short stretches of aberrant amino acids after the frameshift (brown). **(H)** Alignment of wild-type and predicted mutant proteins in the region of frameshift, showing normal sequence (black) and aberrant residues (brown). **(I)** Superimposed time-lapse images (from Movie S1) showing fin motion in a wild-type embryo (yellow crescent arrows) and motionless fins in a *mylpfa*^*oz30*^ mutant (yellow dots) at 4 dpf. Similar results were obtained using a second mutant allele, *mylpfa*^*oz43*^ (not shown). **(J)** Quantification of fin beats per minute averaged over left and right sides, in *mylpfa*^*oz30*^ mutant (n=12) or phenotypically wild-type sibling (n=12) fish at 4 dpf. We have never observed a fin beat in over 100 *mylpfa*^*oz30*^ and *mylpfa*^*oz43*^ mutant fish examined. **(K)** Trunk muscle contractile force in *mylpfa*^*oz30*^ and wild-type or heterozygous siblings at 3 dpf. No significant differences are found between wild-type and heterozygous fish. However, homozygous *mylpfa* mutant fish are significantly weaker than non-mutant siblings at all stimulation frequencies. **(L)** Representative image of fluorescently-labeled actin filaments tracked in the *in vitro* motility assay, with colored lines showing traces of individual filaments over 50 frames of imaging (2.5 seconds). Within this time period, actin filaments on wild-type myosin extracts typically move further than do filaments on *mylpfa* mutant myosin extracts. **(M)** Actin filament speeds measured using the in vitro motility assay generated by extracted myosin. Myosin from *mylpfa* homozygous mutant fish propel actin filaments significantly slower than myosin from wild-type siblings. Numbers shown in each bar indicate the experimental N; each experiment uses myosin extracted from two fish (see Methods). Asterisks indicate P thresholds for the WT/het pools vs mutant; * indicates P<0.05, ** indicates p<0.001, *** indicates p<0.0001. Error bars represent standard deviation. Statistical comparisons in J and M use student’s T test; in K, thresholds are determined by ANOVA followed by Tukey-Kramer comparisons. Scale bar in I is 250 µm.

### Protein analysis

Protein models were downloaded from PDB and visualized using Geneious software. Structures shown have the following PDB accession numbers: scallop IKK7;^40^ Squid 3I5H;^41^ Chicken 2W4G;^42^ Rabbit, 5H53.^43^ Protein alignments were produced by a MUSCLE algorithm in Geneious software. Protein sequences with the following accession numbers were downloaded from NCBI, Ensembl, or Uniprot: human MYLPF (NP_037424.2), *Mus musculus* (mouse) Mylpf (NP_058034.1), *Oryctolagus cuniculus* (rabbit) Mylpf (NP_001076230.2), *Gallus gallus* (chicken) Mylpf (NP_001185673.1), Xenopus tropicalis (frog) Mylpf (CAJ83266.1), *Danio rerio* (zebrafish) Mylpfa (NP_571263.1), *Danio rerio* (zebrafish) Mylpfb NP_001004668.1, *Callorhinchus milii* (elephant shark) skeletal Myl2 (AFP05921.1), *Eptatretus burgeri* (hagfish) Myl2 (ENSEBUT00000005213.1), *Todarodes pacificus* (Japanese flying squid) light chain-2 (LC2; P08052), *Chlamys nipponensis akazara* (Japanese bay scallop) myosin light chain regulatory (MLR; P05963), *Dictyostelium discoideum* (Dicty) RLC (AAA33226.1), and *Saccharomyces cerevisiae* (yeast) Mlc2 (ONH78313.1). The human MYLPF orthologs shown are regulatory light chain genes MLC2 (NP_000423.2), MLC5 isoform 1 (NP_002468.1), MLC7 (AAH27915.1), MLC9 isoform A (NP_006088.2), MLC10 (ENST00000223167.4), MLC12A (NP_001289977.1), and MLC12B (NP_001138417.1) as well as essential light chains MLC1 (NP_524144), MLC3 (NP_524146.1), MLC4 (NP_001002841.1), MLC6 isoform 1 (NP_066299.2), and MLC6 isoform 2 (NP_524147.2). Sequence alignments are ordered by their similarity to human MYLPF. Expression patterns are described previously.^44–46^

## Results

### Identification of variants in MYLPF

Family A, of European ancestry, consisted of two affected siblings born to unaffected parents. Analysis of high-density genotyping data suggested that the siblings were the products of a consanguineous mating (F=0.0189). This observation was later confirmed by review of pedigree records that documented the parents were second cousins once-removed. Each affected child, a male last evaluated at 13 years of age and a female last examined at 24 years of age, had severe contractures of the hands, fingers, wrists, elbows, hips, knees, ankles, and neck; pterygia of the elbows and knees; small mouths; and short stature (< 1 percentile for weight and at ∼1.1 percentile for height (Table 1). Family B comprised an adopted child of East Indian ancestry whose parents were predicted to be first cousins based on analysis of high-density genotyping data (F=0.0657) and who was homozygous for the same variant as found in the siblings in Family A. The proband of Family B, last examined at 6 years of age, also had similar clinical findings to the siblings in Family A including a small mouth, micrognathia, scoliosis, contractures of the hands and wrists, and severe right clubfoot that was recalcitrant to treatment, ultimately leading to amputation of the lower leg. Pathological exam of the foot revealed complete absence of skeletal muscle that was confirmed histologically (Figure 1). In both families, we identified homozygosity for a variant (c.470G>T) in a single gene, *myosin light chain, phosphorylatable fast skeletal muscle* (*MYLPF* [MIM 617378; Refseq accession number NM_013292.4]) (Table 1; Figure 2). This variant is predicted to result in p.Cys157Phe substitution and has a CADD (v1.6) score of 27.5. *MYLPF* had been considered *a priori* to be a high-priority DA candidate gene because of its role in development of skeletal muscle.^10^ Sanger sequencing validated homozygosity of the c.470G>T variant in all affected persons and that the parents in Family A were heterozygous carriers. These results suggested that homozygosity for c.470G>T in *MYLPF* resulted in a pattern of multiple congenital contractures virtually indistinguishable from that observed in persons with DA, specifically DA1.

**Figure 2:**
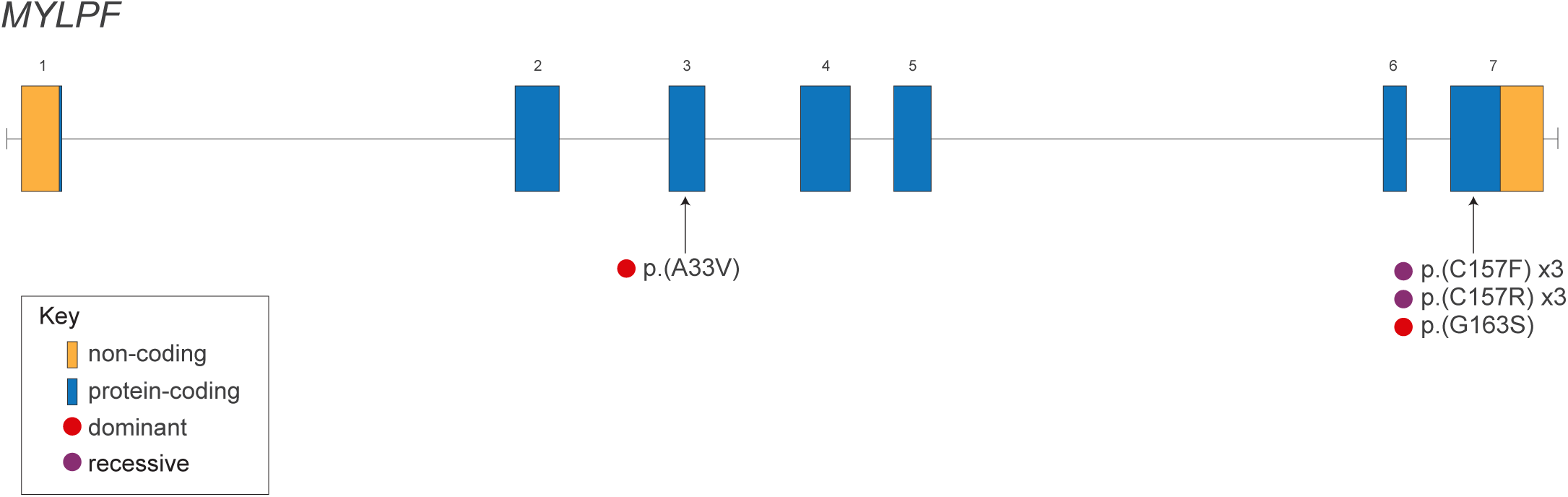
Genomic Model of *MYLPF*. *MYLPF* is composed of 7 exons, each of which consists of protein-coding (blue) and non-coding (orange) sequence. The approximate location of each pathogenic variant is indicated by an arrow. The p.(C157F) and p.(C157R) variants are each found in three families (x3) and lead to a recessive phenotype (purple circle). The p.(A33V) and p.(G163S) variants lead to a dominant phenotype (red circle).

In an effort to find additional families with a pathogenic variant in *MYLPF* and clinical characteristics of DA1, we submitted genetic data and phenotypic information to the MatchMaker Exchange (MME) via the MyGene2 node and identified two additional families (E and F) that had been submitted to the GeneMatcher node. Simultaneously, we queried commercial genetic testing companies and colleagues about families in which *MYLPF* had been identified as a candidate gene. One commercial lab, GeneDx, responded that clinicians for two families (C and D) had agreed to be contacted by us and provide de-identified genetic and phenotypic information for review.

The proband of Family C was a female, born in Pakistan to unaffected parents, and last evaluated at 29 years of age. She had short stature, nearsightedness, mild conductive hearing loss diagnosed in adulthood, scoliosis, bilateral clubfeet, and multiple congenital contractures including the hands, wrists, elbows, shoulders. ES demonstrated that she was homozygous for the variant, c.469T>C (p.Cys157Arg) in *MYLPF*, with a CADD score of 25.2. Family D consisted of an affected stillborn female born to unaffected, first cousin parents from Pakistan. At 35 weeks estimated gestational age a prenatal ultrasound demonstrated polyhydramnios, shortened long bones, “clenched hands” and bilateral clubfeet. The fetus was subsequently stillborn and postmortem examination confirmed the prenatal findings. Proband-only ES of the fetus demonstrated she was also homozygous for the c.469T>C in *MYLPF*. The proband of Family E was a male of Pakistani ancestry, whose parents were first cousins. At ∼13 weeks gestational age, a prenatal ultrasound detected nuchal edema, a septated cystic hygroma, mild enlarged renal pelvis, and normal amniotic fluid. At 19 weeks gestational age, an ultrasound showed decreased fetal movements with fetal hypokinesia and akinesia of lower limbs, clubfeet, thoracic kyphosis and scoliosis, generalized edema (trunk and nuchal) and normal amniotic fluid volume. He had multiple congenital contractures of the hands, wrists, elbows, shoulders, hips, knees and bilateral clubfeet as well as pterygia of the neck. He died at one month of age and was found to be homozygous for the c.469T>C variant in *MYLPF* found in Families C and D. Both parents were heterozygous.

Family F (Table 1 and Figure S1) is a large kindred from South India in which four individuals were each homozygous for c.470G>T in *MYLPF*. Individuals III-2 and III-3 were the offspring of a consanguineous mating between II-1 and II-2 whereas individuals III-5 and III-6, who had two affected fetuses (IV-8 and IV-10), were not known to be closely related but grew up in the same community. Individual III-3 and both affected fetuses (IV-8 and IV-10) had contractures of the hands and feet, while III-2 had contractures of only the hands. None of the heterozygous carriers tested (II-1, II-2, III-5, and III-6) had congenital contractures.

Collectively, we identified two unique missense variants affecting the same residue, p.(Cys157), in *MYLPF* in ten individuals in six unrelated families (A-F) who had been diagnosed with multiple congenital contractures. Both missense variants are exceedingly rare (maximum frequency in any super-population in gnomAD = 0.00013 in South Asians for c.470G>T and 0.00009 in Finnish for c.469T>C, and no homozygotes reported) among >151,000 individuals included in publicly available databases: 1000 Genomes phase 3, the gnomAD browser (v2.0.2), or UK10K (February 15, 2016 release) and have high Phred-scaled CADD scores, consistent with pathogenic dominant variants (Table 1). High-density chip genotype data (Illumina Human Core Exome) was available for one affected individual of East Indian ancestry (Family B) and one affected individual in the family of Polish ancestry (Family A), both of whom were homozygous for the first variant, c.470G>T. We searched for a shared haplotype that would provide evidence that this variant is a founder mutation, but their genotypes differed at even the nearest SNP flanking the variant (both C/C at chr16:30368510 [rs13335932]; both homozygous for c.470G>T at chr16:30389181; but at chr16:30393147 [rs34518080], the Polish individual was A/A while the East Indian individual was C/C), leaving at most a short shared haplotype. Combined with the observation of this variant in multiple populations in gnomAD, it seems likely that this variant has either arisen independently in different populations or early in human history. In contrast, the variant shared among all three Pakistani families, c.469T>C, may be a founder mutation based on their shared ancestry, but we were unable to obtain the original exome sequence data to confirm the presence of a shared haplotype.

In a seventh family (Family G), a one-year-old male proband was found to be heterozygous for a c.487G>A, p.(Gly163Ser) variant that arose *de novo* in his father. The family was of Ashkenazi Jewish origin. The proband, last evaluated at 12 months of age, had multiple congenital contractures including bilateral camptodactyly and overriding fingers, adducted thumbs, ulnar deviation of the wrists, bilateral hip dislocations, and bilateral vertical talus. He had a mild kyphosis, bilateral inguinal hernias, small palpebral fissures, epicanthal folds, anteverted nares with hypoplastic alae nasae, long philtral folds, a thin upper lip, small mouth, high-arched palate without cleft, and micro-retrognathia. Because of recurrent apnea, he underwent a tracheostomy. His father had similar facial features, ulnar deviation of the hands, and had undergone multiple corrective procedures for contractures of the ankles. Manual review in IGV of the proband’s exome sequence data did not reveal any additional rare variants in *MYLPF* and no candidate variants were found in other genes known to underlie DA. Depth of sequencing of all exons of these genes was >10X and there was no evidence of a copy number variant that would explain his features. This variant had a CADD score of 32.0.

Family H is a family in which the two probands, individuals IV-1 and IV-3, were independently diagnosed with distal arthrogryposis (DA). Individual IV-3 was first evaluated at 7 months of age because of a bilateral ulnar finger deviation, flexed thumbs, right calcaneovalgus deformity and left clubfoot. Several months later, her cousin (IV-1) was referred to an arthrogryposis clinic for evaluation of bilateral ulnar deviation and bilateral clubfoot (Figure 1). The mother and grandmother of each proband also had congenital contractures (Table 1). Individual IV-3, last evaluated at 6 years of age, has no growth retardation or scoliosis, neither her mother nor grand-mother. Clinical exome sequencing of IV-3 and III-2 revealed heterozygosity for a c.98C>T p.(Ala33Val) (NM_013292) and Sanger sequencing confirmed it was present in III-6 and II-7 (Table 1 and Supplementary Figure 1). This variant is predicted to be damaging with a CADD score of 31.0 and is absent from gnomAD v2.1.1; accessed on 04/12/2020. Dominant inheritance of DA in Families G and H suggest that p.(Gly163Ser) and p.(Ala33Val) may impact MYLPF function more severely than the recessive variants, p.(Cys157Phe) and p.(Cys157Ser).

### Loss of zebrafish mylpf function causes muscle weakness and loss

Mylpf structure is highly conserved among vertebrates suggesting that function is also conserved. Previous studies showed that mice homozygous for a null *mylpf* allele lack all skeletal muscle at birth.^10^ However, these studies did not determine whether muscle failed to develop or underwent degeneration after normal development, nor did they explain how complete or partial loss of function of MYLPF function in humans selectively and/or disproportionately affects muscles of the limb. To generate a vertebrate model for partial loss of function, we knocked out zebrafish *mylpfa*, one of the two zebrafish *Mylpf* orthologs. Both orthologs, *mylpfa* and *mylpfb*, are expressed specifically in embryonic and larval fast-twitch myofibers,^47^ with *mylpfa* being the predominantly expressed gene (Figure 3A-E). We used CRISPR/Cas9-mediated mutagenesis to generate two independent *mylpfa* alleles, *mylpfa*^*oz30*^ and *mylpfa*^*oz43*^ (Figure 3F). Both alleles frameshift the protein within the first of two EF-hand domains (Figure 3G, H) and are thus predicted to be nulls. However, homozygous *mylpfa* mutant embryos are expected to retain some Mylpf function because they still have a functional *mylpfb* gene.

Pectoral fins of the *mylpfa* mutant are completely paralyzed, and the mutant has an impaired escape response (Figure 3I, J, Movies S1 and S2). To directly measure contractile strength of trunk muscle, we electrically stimulated live intact 3 dpf embryos mounted between a hook at the head and a force transducer at the tail. At all frequencies tested, *mylpfa*^*oz30*^ homozygous mutant embryos are significantly weaker than their unaffected siblings (P<0.001 at 20 Hz and 180 Hz) (Figure 3K). To characterize the motion-generating capacity of myosin with their constituent light chains, we extracted monomeric myosin from *mylpfa* mutant and sibling wild-type fish onto the surface of a microscope flow cell chamber (see Methods). Fluorescent, filamentous actin was introduced into the chamber and actin motion generated by the extracted myosin was imaged. We observed that the actin filament velocity was significantly slower in both *mylpfa*^*oz30*^ and *mylpfa*^*oz43*^ compared to wild-type siblings (Figure 3L, M, Movie S3). The remaining and apparently compromised myosin motile function and contractile force observed could be due to residual *mylpf* function in fast-twitch muscles (provided by *mylpfb*) and/or intact slow-twitch muscles, which do not express *mylpfa* and appear normal (Figures 3A-D and 4C-F). Thus, like persons with DA, zebrafish *mylpfa* mutants display abnormal movement that is more pronounced in limbs; this weakness can be explained at least in part by impaired myosin force and motion generation in fast-twitch muscle.

We next examined zebrafish muscle over time, to learn whether *mylpfa* mutant muscles form and then deteriorate or whether they fail to develop in the first place. This distinction may be particularly important for understanding the complete absence of skeletal muscle that was observed in the foot of one child with DA associated with a p.(Cys157Phe variant). Shortly after somite formation, *mylpfa* mutant muscle morphology appears normal (Figure 4A, B). However, at 6 days post fertilization (dpf), *mylpfa* mutant myotomes are significantly (p<0.05) reduced in dorsal-ventral height compared to wild-type (WT = 266 µm, N = 12; *mylpfa*^*-/-*^=232 µm, N=6) and muscle fibers are irregularly shaped, suggesting that fast-twitch fibers deteriorate over time (Figure 4C, D). Slow-twitch muscle fibers are spared in the same fish (Figure 4E, F). Muscle defects are most pronounced in a specific appendicular muscle, the posterior hypaxial muscle (PHM) (Figure 4G-J). Although the PHM forms at the normal time in *mylpfa* mutants and is initially comprised of multinucleate muscle, the muscle fibers break down into small ‘islands’ that are often mononucleate by 4 dpf (Figure 4I, J). Muscle deterioration in the PHM is especially rapid compared to axial muscle, since axial muscle fibers form prior to appendicular fibers, yet are still relatively intact and multinucleate at 6 dpf (compare Figure 4D and J insets).

**Figure 4:**
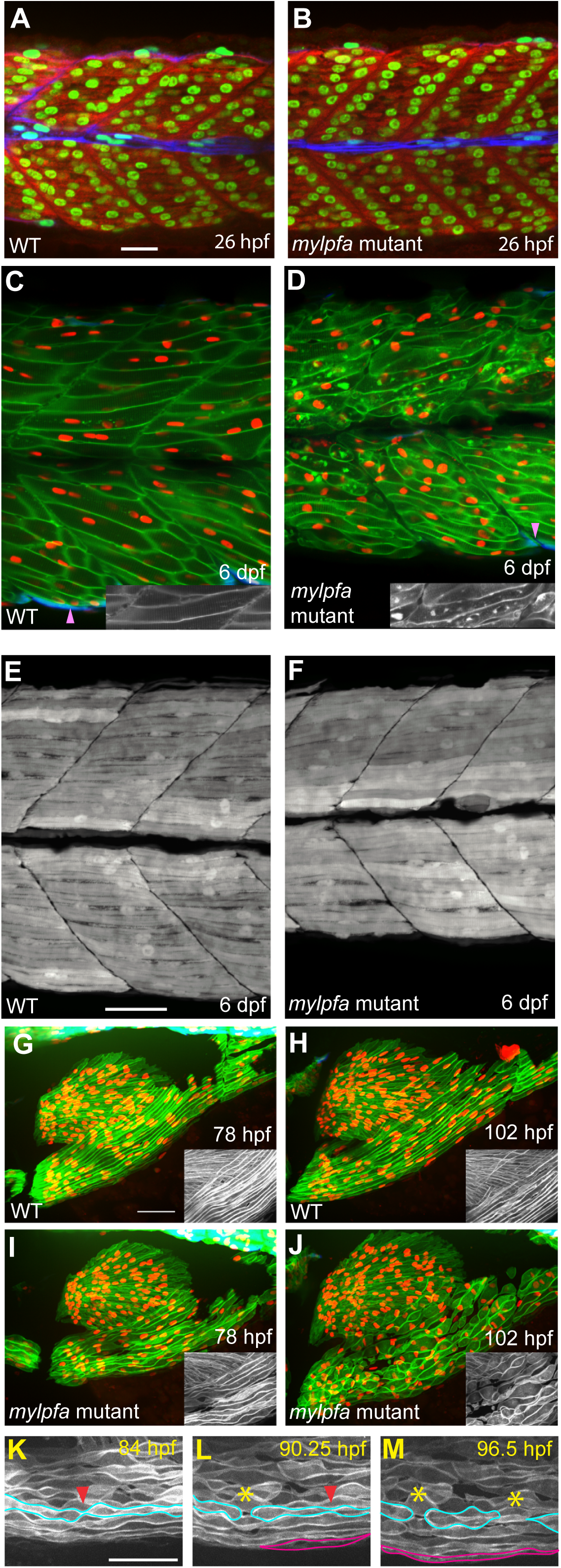
Embryonic muscle degenerates in *mylpfa* mutant zebrafish. **(A, B)** Confocal images showing muscle morphology at 26 hpf in wild-type (A) or *mylpfa*^*-/-*^ (B) embryos. Myonuclei are labeled using Rbfox1l immunolabeling (green), fast-twitch fibers are labeled using F310 immunolabeling (red), and slow-twitch fibers are labeled using *Tg(smyhc1:EGFP)i104* (blue). **(C, D)** Confocal z-sections showing muscle morphology of live 6 dpf larvae expressing a fast muscle cell membrane transgene *Tg(mylpfa:lyn-Cyan)fb122* (green) and a myonuclear transgene *myog:H2B-mRFP* (red). These fish also express a slow muscle marker *Tg(smyhc1:EGFP)i104* (blue), which is largely lateral to the plane of focus; pink arrowheads point to slow muscle cells within the shown plane. Compared to wild-type controls (C), *mylpfa*^*-/-*^ myofibers have irregular membrane structure (D). **(E, F)** Confocal projections of the same sections from panels C and D showing slow muscle fibers (white). **(G-J)** Confocal projections of pectoral fin and PHM muscles, imaged on 3.25 dpf and again on 4.25 dpf, in fish expressing transgenes *Tg(mylpfa:lyn-Cyan)fb122* in fast muscle (green), *myog:H2B-mRFP* in all myonuclei (red), and *Tg(smyhc1:EGFP)i104* in slow muscle (blue). Fin muscle and PHM express *Tg(mylpfa:lyn-Cyan)fb122* but not *Tg(smyhc1:EGFP)i104*, as expected for muscles predominantly comprised of fast-twitch fibers.^23,57^ In striking contrast to wild-type (G, H), *mylpfa* mutant muscle fibers degenerate between 3.25 and 4.25 dpf (I, J). **(K-M)** Images from a time-lapse of *mylpfa* mutant PHM degeneration (Movie S3). Muscle fibers which initially appear wavy (aqua outline), often become narrow (red arrowheads) before breaking apart (asterisk). A myofiber that appears during imaging is outlined in magenta. Insets in C, D, G-J show *Tg(mylpfa:lyn-Cyan)fb122* in greyscale. Scale bars in A (for A, B), E (for C-F), G (for G-J), and K (for K-M) are 50 µm.

To examine whether *mylpfa* mutant PHM cellular ‘islands’ are degenerated fibers or newly forming mononucleate myoblasts, we conducted confocal time-lapse microscopy beginning at 84 hpf (Figure 4K-M; Movie S4). At time-lapse outset, most PHM fibers are intact, with wavy membranes that occasionally are narrowed almost to closure (Figure 4K, L). Membrane irregularities become more pronounced over time, and pieces of fiber pinch off (Figure 4L, M). During each time-lapse (N=3), a new PHM fiber was added (Figure 4L, M), suggesting that hyperplastic growth continued during the imaging period. Together, these findings indicate that zebrafish *mylpfa* is required to maintain myofiber integrity but is not required for initial myofiber formation or hyperplastic growth. Accordingly, we hypothesize that the segmental amyoplasia in individuals with pathogenic variants results myofibers that form normally but subsequently degenerate.

### *Protein modelling of* MYLPF *variants*

To better understand how MYLPF p.(Cys157Phe) and p.(Cys157Arg) variants versus the p.(Ala33Val) and p.(Gly163Ser) variants underlie recessive versus dominantly inherited DA, respectively, we examined a previously developed protein model of the rabbit Mylpf-Myl1-Myosin heavy chain-Actin complex (PBD 5H53)^43^ (Figure 5A, B). Rabbit Mylpf protein is shifted one amino acid relative to the corresponding residues in the human counterparts (Figure 5C) but for simplicity, we refer to equivalent MYLPF/Mylpf residues using human numbering across species. Ala33 is conserved in all eukaryotes examined, Gly163 is deeply conserved from scallop to human, but Cys157 is conserved only among vertebrates (Figure 5C). The protein model reveals that Ala33 is positioned adjacent to the myosin heavy chain and Mylpf Gly163 directly contacts a phenylalanine near the hook region of myosin heavy chain (Figure 5A). In contrast, Mylpf Cys157 is buried deep within the regulatory light chain and makes no contact with myosin heavy chain (Figure 5A).

**Figure 5:**
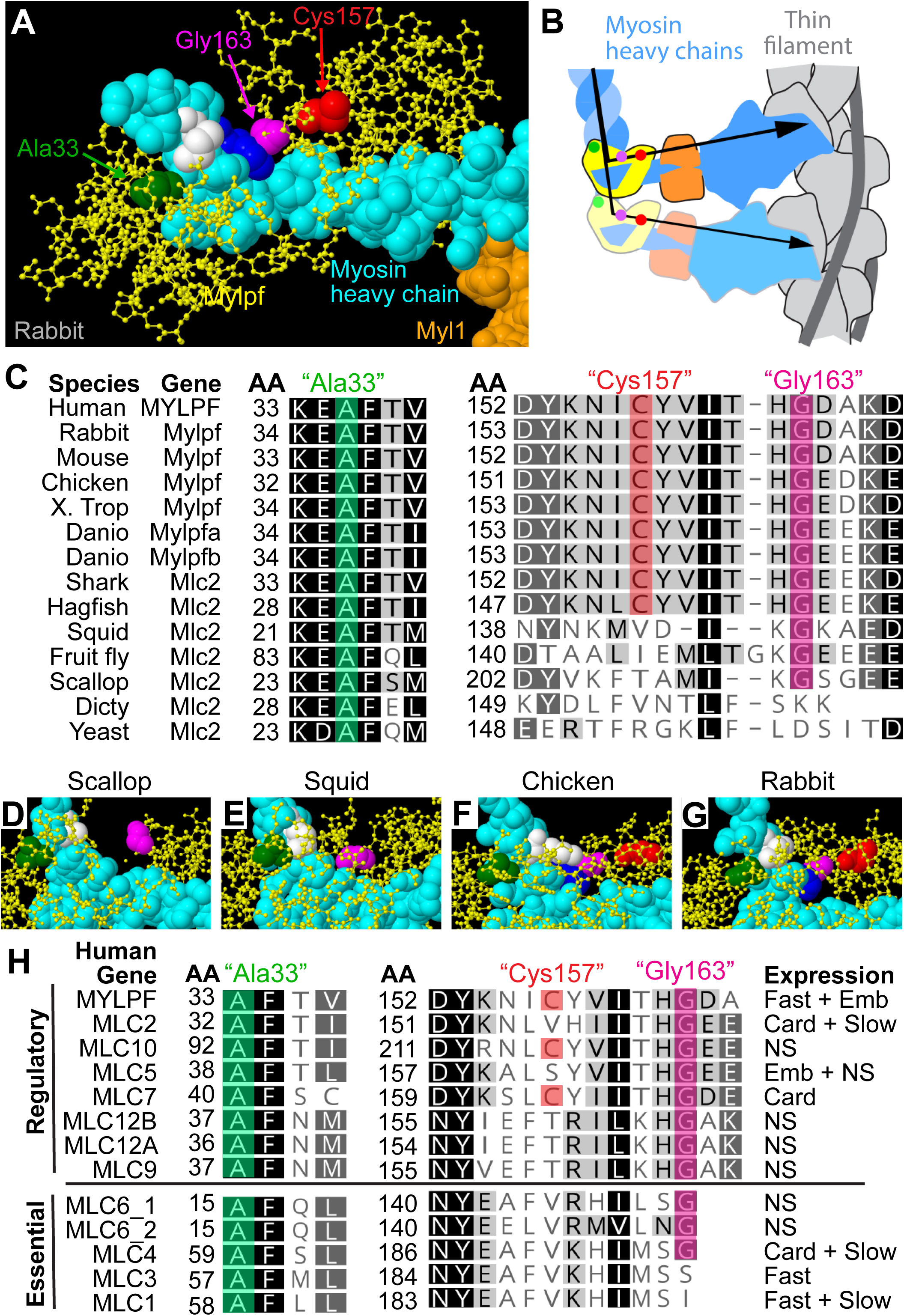
Mylpf protein sequence and structure comparisons identify key conserved residues. **(A)** Model of rabbit Mylpf protein in complex with the neck and head region of myosin heavy chain in rigor. The heavy chain (cyan) and essential light chain (orange) are rendered using a space-filling model and the light chain is shown using a ball and stick model (yellow) except for three residues that align with MYLPF Ala33, Cys157, and Gly163, which are rendered in space filling models; we refer to these by their human numbering. Ala33 (green) is adjacent to a Lys residue in the heavy chain (white). Gly163 (magenta) directly contacts a Phe residue in the heavy chain (dark blue). In contrast, Cys157 (Red) is found internal to Mylpf protein. **(B)** Overview of myosin interaction with thin filaments, color-coded as in panel A. Mylpf protein binds to the heavy chain region that bends towards the thin filament (arrows). **(C)** Alignment of select vertebrate Mylpf proteins and invertebrate Myl2 proteins highlighting conservation of Ala33, Cys157, and Gly163. **(D-G)** Magnified views of myosin heavy and light chain genes showing how Ala33 and Gly163 positions vary between scallop^40^ (**D**), squid^41^ (**E**), chicken^42^ **(F)**, and rabbit^43^ **(G)**. Cys157 is located internally to the two vertebrate Mylpf structures (F, G). Color coding in panels D-G is the same as in panel A. **(H)** Alignment of human regulatory and essential light chain proteins highlighting conservation of Ala33, Gly163, and Cys157. The tissue that express each ortholog is indicated as follows: embryonic skeletal muscle (Emb), fast-twitch skeletal muscle (Fast), slow-twitch skeletal muscle (Slow), cardiac muscle (Card), and non-sarcomeric tissue (NS). The first residue in the shown aligned portions are numbered for each protein, and proteins are arranged by their similarity to MYLPF (C and H).

To determine whether the interaction between Mylpf Gly163 and myosin heavy chain is conserved, we compared crystal structures of scallop, squid and chicken myosin light and heavy chains to the rabbit structure (Figure 5D-G; structures generated in^39–42^). In chicken, as in rabbit, Gly163 directly contacts a phenylalanine residue in the heavy chain (Figure 5F, G). Gly163 is found on the surface of the RLC in scallop and positioned very close the myosin heavy chain in squid (Figure 5D, E). Because Gly163 is present in animals that diverged from vertebrates prior to regulatory light chain gene family diversification, we reasoned that MYLPF orthologs may also contain this residue. Alignment of human MYLPF orthologs reveals perfect conservation of glycine in the position corresponding to human Gly163. Some essential light chain proteins also have a corresponding glycine, suggesting that it arose very early in light chain gene evolution (Figure 5H). Likewise, Ala33 is conserved in all light chains examined including both ELCs and RLCs and is positioned close to the heavy chain in the crystal structures examined (Figure 5C-H). In contrast, only three of the eight human regulatory light chain genes contain a cysteine in the Cys157 position (Figure 5H), revealing that although Cys157 is conserved among vertebrate Mylpf genes (Figure 5C), some orthologous regulatory light chain genes have a different residue in this position. Together these findings indicate that Ala33 arose prior to animal evolution, Gly163 arose early in animal evolution, and these residues directly or almost-directly contact the myosin heavy chain in vertebrates. These observations suggest that the *MYLPF* p.(Ala33Val) and p.(Gly163Ser) variants have dominant effects because they more directly impact interactions with myosin heavy chain than the Cys157 variants.

## Discussion

We identified four pathogenic variants in *MYLPF* in eight unrelated families in which seventeen affected individuals share similar phenotypic features, suggesting that mutations in *MYLPF* underlie a previously unrecognized multiple malformation syndrome. This condition is characterized by multiple congenital contractures, scoliosis, and short stature and is clinically indistinguishable from DA1 due to mutations in genes encoding other components of the contractile apparatus (e.g., embryonic myosin, troponins, tropomyosin). However, we find that short stature and proximal joint contractures (i.e., elbows, hips, knees) appear more commonly in DA1 due to variants in *MYLPF* than in DA1 due to other variants in other genes (*MYH3, TNNI2, TNNT3* and *TPM2*).^5,6^ Nevertheless, because the number of families with pathogenic variants reported is small, the extent of overlap between their phenotypic features and those of individuals with variants in *MYH3, TNNI2, TPM2* and *TNNT3* remains to be determined.

In contrast to the clinical findings of DA1 specifically, and of DAs in general, a single individual had segmental amyoplasia of the foot with fatty replacement of the muscle. We did not observe this finding in other persons with DA due to *MYLPF* variants, but we lacked pathological or imaging data of the limbs for all but one additional case, an affected fetus that underwent post-mortem exam at 19 weeks. Whether this is a finding in other individuals with pathogenic variants in *MYLPF* is unclear. To our knowledge, complete absence of limb skeletal muscles in a person with DA has not been reported. Hypoplasia or aplasia of the muscles of the upper and / or lower limbs is the defining feature of a group of arthrogryposis conditions known as Amyoplasia.

Amyoplasia is the most common condition referred to as arthrogryposis, accounting for ∼30% of all persons diagnosed with arthrogryposis.^48^ The etiology is suspected to be heterogeneous (i.e., vascular disruption, monogenic, somatic mosaicism, oligogenic, etc.) and the heritability of Amyoplasia, if any, remains unknown. The overwhelming majority of cases of Amyoplasia are simplex, but rare instances of affected siblings have been reported. In such cases, amyoplasia is typically limited to either the upper or lower limbs.^49^ In a small subset of such families with lower limb amyoplasia (LLA), pathogenic variants have been found in one of several genes including *BICD2*,^50^ *CACNA1H*,^51^ *DYNC1H1*,^52^ *TRPV4*,^53^ and *FKBP10*.^54^ In each of the LLA conditions resulting from pathogenic variants in these genes, neurological abnormalities including weakness and hypotonia are typically present, distinguishing them from DA1 due to *MYLPF* variants. However, absence or severe atrophy of select muscles of the lower limbs is also common, if not typical, in these conditions, suggesting segmental amyoplasia is a genetically heterogeneous trait. Moreover, these observations suggest that, at least in some families with Amyoplasia, large effect risk allele(s) might be segregating.

Our findings indicate that DA1 due to pathogenic variants in *MYLPF* can be transmitted as either an autosomal dominant or autosomal recessive condition. Nearly 400 genes underlying Mendelian conditions transmitted in both dominant and recessive inheritance patterns have been reported (J.X. Chong et al., 2019, Am. Soc. Hum. Genet., abstract). In the majority of these instances, the resulting Mendelian conditions have overlapping but different phenotypic features suggesting that while the inheritance patterns may differ, the pathogenesis for each is similar. In some cases, including DA1 due to variants in *MYLPF*, the clinical characteristics of the dominant and recessive conditions are virtually indistinguishable (e.g., cataracts due to variants in *CRYAA*). Differences in inheritance patterns can result from a variety of phenomena, including variants that affect distinct functional domains, result in loss versus gain-of-function, affect different tissue-specific transcripts, and have dose-dependent effects on gene function.

Substitutions of Gly163 and Ala33 in MYLPF both result in dominant DA1 whereas substitution of Cys157 underlies recessive DA1. These three residues are each conserved among all vertebrate MYLPF homologs, but Ala33 and Gly163 are more deeply conserved than Cys157. Indeed, Ala33 is found in all light chain genes examined and Gly163 is found in all animal RLC genes reviewed. This broad conservation suggests that Ala333 and Gly163 are vital to RLC identity. These two residues sit on the surface of MYLPF that contacts MyHC, whereas Cys157 is buried within MYLPF. This indicates that Ala33 and Gly163 might directly participate in critical protein-protein interactions. Perturbation of these residues may directly affect the packing of MyHC molecules into the thick filament, which might in turn alter the function of other nearby myosins in this multimeric structure. Disruption of MYLPF amino acid residues that are positioned further from MyHC (e.g., Cys157) may result in lesser adverse effects on force transduction.

Based on the rabbit crystal structure, Ala33 and Gly163 may contact residues in human embryonic myosin, MYH3, that are affected in other DA conditions. Specifically, Gly163 is predicted to directly contact a phenylalanine residue in the MyHC that corresponds to Phe835 in MYH3. We previously reported an alteration at this MYH3 residue (c.2503_2505delTTC; p.(Phe835del)) in an individual with DA2A, the most severe of DA conditions. Similarly, Ala33 is positioned directly adjacent to a residue in MyHC that corresponds to MYH3 Lys838, a residue that we previously reported to be altered (c.2512A>G; p.(Lys838Glu)) in an individual with DA2B. In both cases, the condition (DA1) resulting from perturbation of MYLPF is less severe than the conditions (DA2A and DA2B) due to altering corresponding residues in MYH3. This difference in disease severity may reflect differences in protein function as MYLPF is thought to act primarily by stabilizing MyHC structure.^7,55^ We predict that other deeply conserved MYLPF residues that are positioned adjacent to disease-associated MyHC residues, such as MYLPF residues Glu29 and Glu32, may, if altered, also lead to dominantly inherited DA1.

We showed that myosin extracted from *mylpfa*-deficient zebrafish moves actin filaments more slowly than wild-type myosin extracts. The slowing (∼75% of full speed) is less pronounced than was found in previous studies which removed myosin RLCs *in vitro* (∼33% of full speed)^8,9^ and compared to *in vitro* extracted chicken myosin bearing a point mutation in Mylpf p.(Phe102Leu) (∼50% of full speed). ^56^ We speculate that the effect of *mylpfa* loss is milder than was seen in these previous assays because the mutant still has some RLC function, provided by *mylpfb* in fast fibers and by a normal set of RLCs in slow-twitch fibers. While the effect on actin motility is modest, total trunk muscle force is dramatically decreased in *mylpfa* mutants and the pectoral fins are completely paralyzed. This parallels the phenotypic consequences observed in DA1, and DAs in general, in which the more distal body areas (e.g., hands vs. shoulders) are more frequently and more severely affected.

In addition to muscle weakness and pectoral fin paralysis, zebrafish *mylpfa* mutant muscle fibers eventually degenerate in all muscles, with the PHM being most severely affected. This degeneration indicates that Mylpf function is essential for muscle integrity during early development and suggests mechanisms for muscle loss in humans with DA1. *Mylpfa* is a marker of fast-twitch myofibers and is abundantly expressed in PHM whereas there are few *smyhc2*-positive slow twitch fibers and no *smyhc1*-positive slow-twitch fibers in the PHM suggesting that it is composed largely of fast-twitch myofibers.^23,30,57^ In contrast, slow-twitch myofibers are more common in myotomes where they may exert a stabilizing effect. Our finding that the PHM is most strongly affected is consistent with the observation that the most severe contractures in DA1, as well as muscle hypoplasia and/or aplasia, occur more frequently in the most distal regions of the limb (e.g., digits, hands, wrists, feet, and ankles). Muscle fibers in myotomes also become irregularly shaped over time in *mylpfa* knockouts, consistent with the observation that the trunk muscles of persons with DA1 due to *MYLPF* variants are typically affected but more mildly than the limbs.

In summary, we discovered that variants in *MYLPF* underlie both dominant and recessive forms of a distal arthrogryposis with features typically seen in DA1. The distribution of these features in persons with DA1 due to *MYLPF* variants largely overlaps that of DA1 due to variants in *MYH3, TNNI2, TPM2*, and *TNNT3*, with proximal joint contractures perhaps more common in individuals with *MYLPF* variants. However, segmental amyoplasia appears to be a unique feature of DA1 due to *MYLPF* that, based on knockout of *mylpfa* in zebrafish, results from degeneration of differentiated skeletal myofibers.

## Supplemental Materials and Methods

Supplemental information includes one figure (S1), four movies (S1-S4), and supplemental methods.

## Acknowledgements

We thank the families for their participation and support. Sequencing was provided by the University of Washington Center for Mendelian Genomics (UW-CMG) and was funded by NHGRI and NHLBI grants UM1 HG006493 and U24 HG008956, by the Office of the Director, NIH under Award Number S10OD021553, and by the National Institute of Child Health and Human Development (1R01HD048895 to M.J.B.). We thank the Ohio State Rightmire Hall zebrafish staff for excellent animal care and husbandry and Mark Nilan and Mika Gallati for zebrafish care and mutant identification at the University of Maine. We thank Sarah Shepherd for assistance during initial *mylpfa*^*oz43*^ construction. Zebrafish work was funded by NIH grants GM088041 and GM117964 (to S.L.A.), an NIH T32 training grant NS077984 and an Ohio State Pelotonia postdoctoral fellowship (to J.C.T.), Ohio State Edward Mayers and Elizabeth Wagner research scholarships (to E.M.T), and HL150953 and AR067279 (to D.M.W.). The Ohio State Neuroscience Imaging Core facilities are supported by NIH grants P30-NS045758, P30-NS104177 and S10-OD010383. We acknowledge the DNA Sequencing Shared Resource at The Ohio State University Comprehensive Cancer Center. This work was also supported by the Indian Council of Medical Research, Government of India (No.5/13/58/2015/NCD-III). The F310 antibody developed by Frank Stockdale was obtained from the Developmental Studies Hybridoma Bank, created by the NICHD of the NIH and maintained at The University of Iowa, Department of Biology. The authors are grateful to Mrs Séverine Drouhin and to the Molecular Biology Facility of the Grenoble University Hospital, France for technical assistance. The content is solely the responsibility of the authors and does not necessarily represent the official views of the National Institutes of Health.

## Web resources

The URLs for data presented herein are as follows:

Geno2MP: https://geno2mp.gs.washington.edu/Geno2MP/

gnomAD: http://gnomad.broadinstitute.org

Human Genome Variation: http://www.hgvs.org/mutnomen/

Online Mendelian Inheritance in Man (OMIM): http://www.omim.org/

PDB: http://www.rcsb.org/

EMBL-EBI expression atlas: https://www.ebi.ac.uk/gxa/home

