## Supplemental Methods and Legends for "Mutations in *MYLPF* cause a novel segmental amyoplasia that manifests as distal arthrogryposis"

**Supplemental Materials and Methods**

**Movie S1: Pectoral fin paralysis in the zebrafish *mylpfa* mutant. (A)** 1-minute video of a wild-type sibling larva showing normal pectoral fin movements. **(B)** 1-minute video of a *mylpfa* mutant, showing that pectoral fins are paralyzed.

**Movie S2: Escape response defects in *mylpfa* mutant zebrafish. (A-B)** Video of wild-type sibling and *mylpfa* mutant larvae showing evoked escape response at 4 dpf. When stimulated, wild-type fish (A) rapidly left the imaging area but *mylpfa* mutants (B) only sometimes escaped and moved slowly even when successful (WT = 12/12 escapes, 0.09±0.02 seconds to exit; *mylpfa* = 7/12 escapes, 4.2±2.2 seconds to exit; $P<0.01$).

**Movie S3: Myosin propels actin filaments more slowly in *mylpfa* mutant extracts compared to wild-type extracts. (A, B)** Video of actin filament movement on myosin extracted from wild-type (A) or *mylpfa* mutant (B) embryos at 4 dpf. For each video, ten actin filaments are tracked as indicated by different colored lines. Over the course of imaging, actin filaments on wild-type myosin extracts tend to move further, suggesting faster filament speeds than for myosin from *mylpfa* mutant fish. Similar results were obtained using fish from *mylpfa*$^{oz30}$ (WT N=10 flow cells; mutant N=8 flow cells) and *mylpfa*$^{oz43}$ (WT N=6 flow cells; mutant N=4 flow cells) crosses.

**Movie S4: The posterior hypaxial muscle degenerates in *mylpfa* mutant zebrafish.** Time-lapse imaging of the PHM in a transgenic embryo expressing *Tg(mylpfa:lyn-Cyan)fb122* (green) and *myog:H2B-mRFP* (red). Images were collected every 5 minutes from 84 to 101 hours post fertilization (hpf). This movie is representative of two time-lapse movies. Scalebar is 50 μm.

**Figure S1. Pedigrees of Families A-H.**

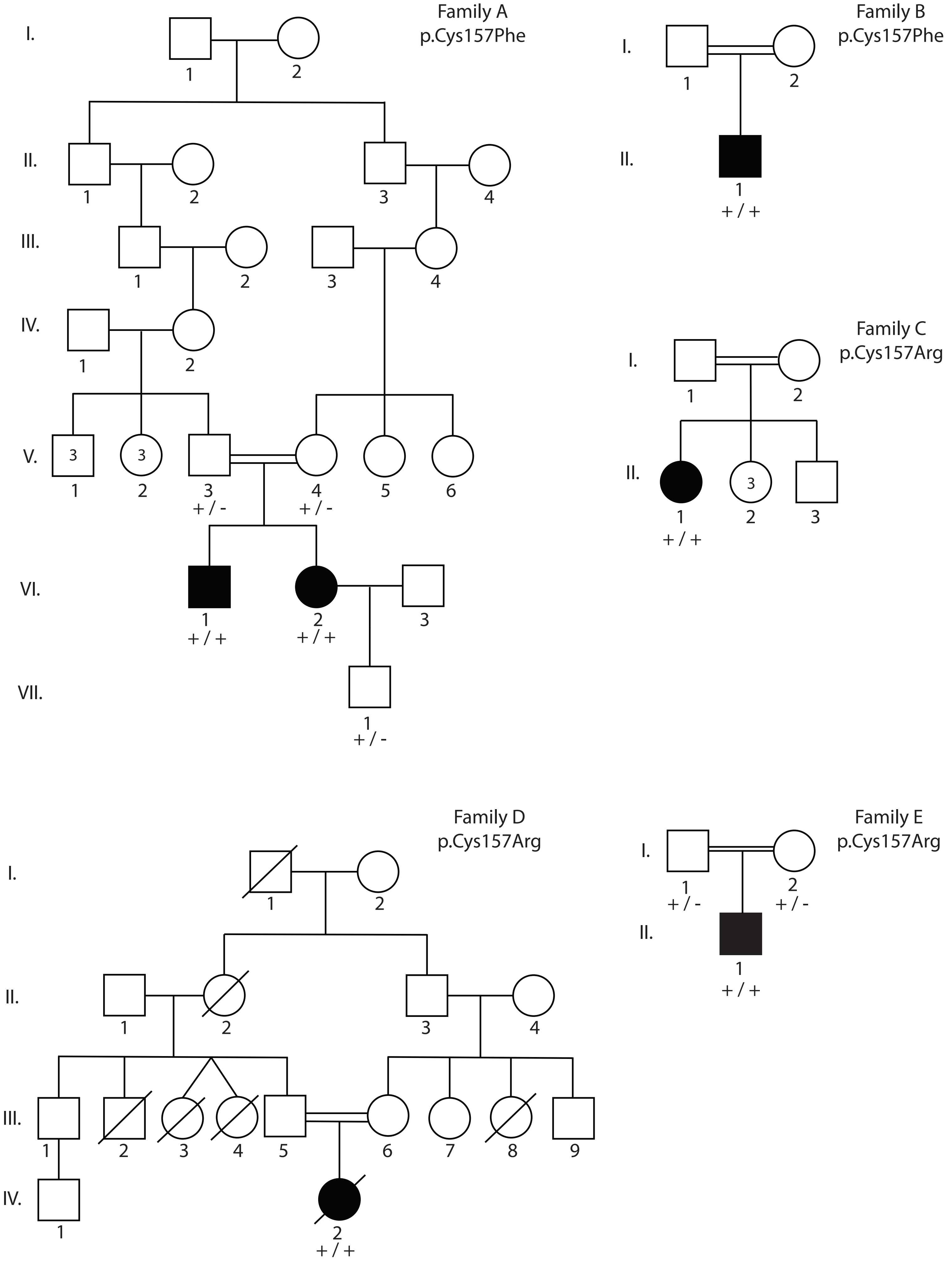
Family A
p.Cys157Phe
I.
II.
III.
IV.
V.
VI.
VII.
+ / -
+ / -
+ / +
+ / +
+ / -
Family B
p.Cys157Phe
+ / +
Family C
p.Cys157Arg
+ / +
Family D
p.Cys157Arg
+ / +
Family E
p.Cys157Arg
+ / -
+ / -
+ / +

Family F
p.Cys157Phe

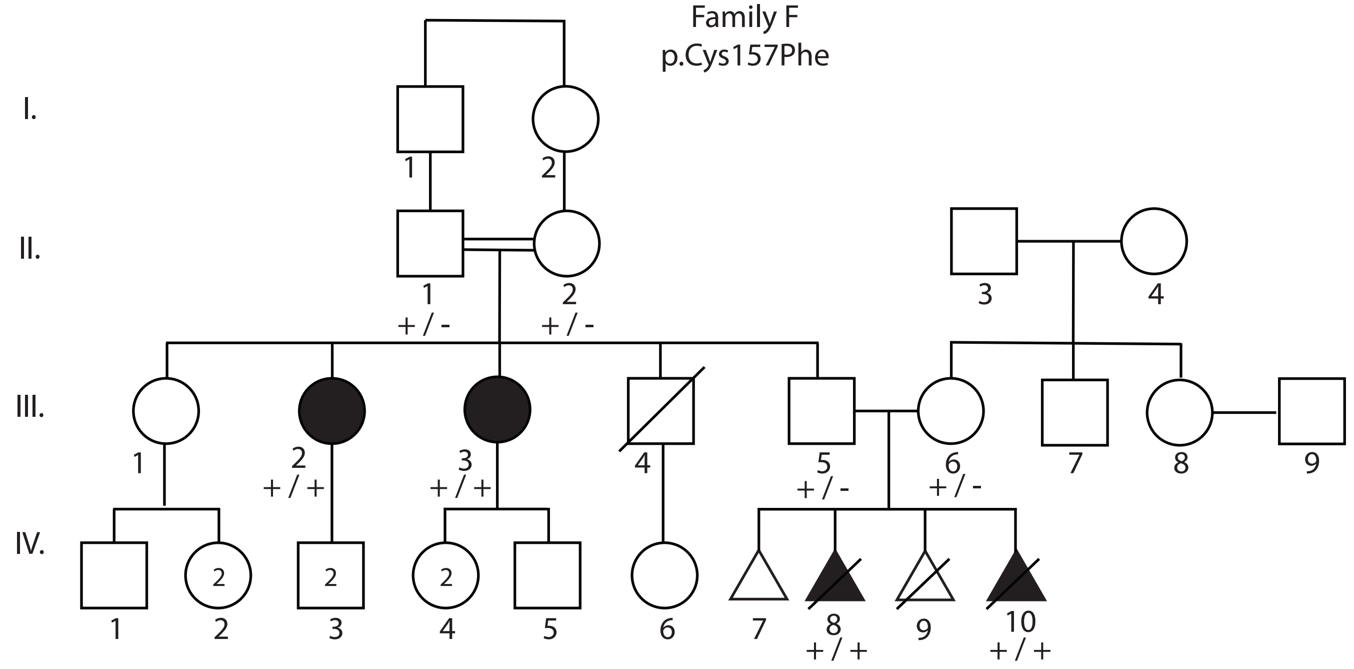


Family G
p.Gly163Ser

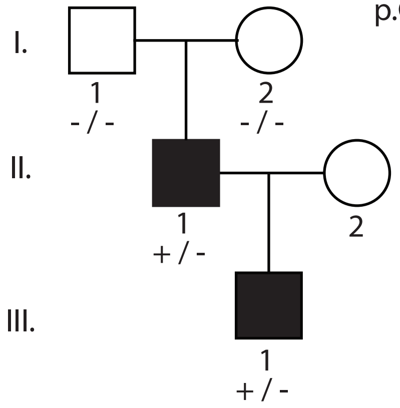


Family H
p.Ala33Val

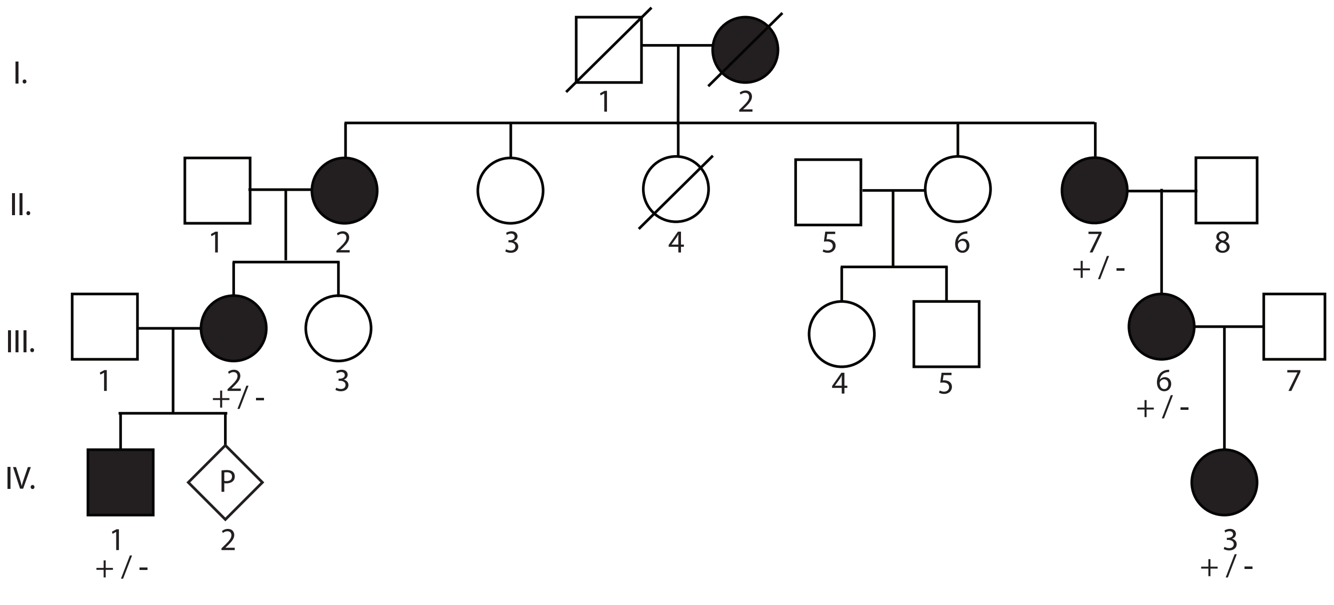

**Methods**

***In Vitro* Motility Assay (IVMA) for zebrafish larvae**

This IVMA procedure was adapted from "***Warshaw et al (1990) J Cell Biol. vol. 111 (2) pp. 453-463"*** and "***Palmiter et al (2000) J Muscle Res Cell Motil. 21(7) pp. 609-20.***" Imaging as described in "***Li et al (2019) Proc Natl Acad Sci U S A. 116(43) pp. 21882-21892.***" Modifications to these protocols are as described below:

**Prepare IVMA flow cells (can be done in advance):**
___ Coat 22mm x 60 mm glass coverslips with nitrocellulose
___ Cut plastic shims of ~ 2 x 30 x 0.13 mm
___ Coat shims with UV glue and apply to nitrocellulose coated coverslips
___ Apply to glass and adjust as necessary to the number of lanes you want
___ Place a plain 22 x 22 mm glass coverslip on top (to form the flow cell chambers)
___ Cure glue under UV light for ~10 minutes

**Prepare embryo for IVMA:**
___ Obtain zebrafish embryos and raise to 4 dpf. Separate fish into phenotypic classes before proceeding further.
___ Dechorionate any unhatched fish using no. 5 watchmakers forceps.
___ De-yolk larvae as follows:
    ___Manually dissect yolks in 2 ml Ringer's lacking $CaCl_2$ plus 200 µl Tricaine
    ___Transfer to a new plate of Ringer's lacking $CaCl_2$
    ___ Wash at least five minutes on a horizontal shaker, to eliminate yolk granules
___ Treat 7.5 minutes in 2 ml protease solution w. 200 µl Tricaine, in a 28.5˚C incubator
    ___ Prewarm the protease and stop solutions prior to use.
___ Add 500 µl Stop Solution
___ Pause 1 minute
___ Transfer the fish to a plate with 2 ml Ringer's plus 200 µl Tricaine
___ Use insect pins to gently open the abdomen and loosen the muscle fibers, to allow the remaining buffers to permeate throughout the fish.
___ Transfer embryos to skinning buffer for 30 minutes. This will permeabilize cells throughout the embryo for extraction. Ideally do this in a petri dish, and swirl a few times.

**Extract myosin from embryo into the flow cell**
___ Transfer two larvae into the flow cell; use a p20 pipetter set to ~5 µl.
___ Using an insect pin, orient the fish so the tail faces into the flow cell, at the center of the flow cell's width. Subsequent washes will pull the tail inwards, while the larvae's wide head and fins will prevent the larvae from being pulled all the way into the flow cell.
___ Add 20 µl of high-salt extraction buffer to skinned embryos for 1 hour on ice at ~40 degree angle. The high ionic strength of this buffer will cause myosin in fibers to depolymerize and flow out of the fish.
___ Place flow cell on a warming plate set at 30°C, tilted to a ~40˚ angle
    *At this point, the slide is coated with monomeric myosin from the larvae.*

**Wash flow cell to eliminate non-functional myosin heads**
*Non-functional myosin heads bind strongly to actin and can serve as a load to freely moving actin filaments. They can be effectively eliminated by applying unlabeled actin, which will be irreversibly bound by the non-functional myosin, prior to beginning the IVMA.*
___ Block surface with a BSA wash (each wash is twice the volume of the flow cell)
___ Coat the flow cell with unlabeled actin for 1 minute
___ Wash 1 x with actin buffer + 1 mM ATP.
___ Wash 1 x with actin buffer

**Apply labeled actin and begin the *in vitro* motility assay.**

___ Add labeled actin buffer to the flow cell for 1 minute
___ Wash 1 x with actin buffer.
___ Activate myosin with motility buffer
___ Transfer flow cell to a microscope stage and oil-couple to a heated objective (30˚ C).
___ Allow the flow cell to temperature equilibrate for 30 seconds to 1 minute
___ Image with 100x objective (n.a. 1.4), 20 frames/second. Image multiple fields of view for each flow cell.
___ During imaging, re-apply motility buffer as needed.

**Solutions:**

**Ringer's solution:** *114 mM NaCl  2.9 mM KCL  5 mM Hepes, pH 7.2*

| | | |
|---|---|---|
| 725 µl 1M KCl | 5.8 ml 5M NaCl | 1.25 ml 1M Hepes, pH 7.2 |
| 242 ml $H_2O$ | Filter sterilize | |

**Protease solution:** *0.25% Trypsin  1 mmM EDTA, pH 8.0  1X PBS*
25 mg Trypsin, Type IV-S (Sigma T0303; Store @ -20˚C before use)

| | | |
|---|---|---|
| 20 µl 0.5M EDTA | 1 ml 10x PBS | 9 ml sterile $H_2O$ |

**Stop solution** *25% fetal calf/bovine serum  500 µM CaCl2   1X PBS*
2.5 ml fetal calf serum (or fetal bovine serum)

| | | |
|---|---|---|
| 50 µl 1M CaCl2 | 1 ml 10x PBS | 6.5 ml sterile $H_2O$ |

**Skinning buffer:** *20mM HEPES (pH 7), 10 mM EGTA, 4.42 mM Mg(OH)2, 8 mM ATP, 10 mM Creatine phosphate, 10 mM DTT, 50% glycerol*

**Extraction buffer:** *150 mM potassium phosphate, 300 mM KCl, 1mM MgCl2, 10 mM Sodium phosphate, 2 mM ATP, 1 mM DTT*

**Actin buffer** *25 mM KCl, 1 mM EGTA, 10 mM DTT, 25 mM Imidazole, 4 mM MgCl2*

**Labeled Actin Buffer:** *Actin buffer, but with Actin filaments labeled with Rhodamine-Phalloidin or another color*

**Motility buffer:** *Actin buffer plus 0.5% methylcellulose, 1 mM ATP, 0.1 µg/ml glucose oxidase, 0.018 µg/ml catalase, 2.3 µg/ml glucose.*
